# Hyper-connectivity between the left motor cortex and prefrontal cortex is associated with the severity of dysfunction of the descending pain modulatory system in fibromyalgia

**DOI:** 10.1101/2021.02.11.430747

**Authors:** Álvaro de Oliveira Franco, Camila Fernandes, Paul Vicunha, Janete Bandeira, Maria Adelia Aratanha, Iraci Lucena da Silva Torres, Felipe Fregni, Wolnei Caumo

## Abstract

The impaired cortical function likely plays a critical role in chronic pain maintenance and fibromyalgia symptoms. Functional connectivity (FC), as assessed by functional near-infrared spectroscopy (fNIRS), is a promising approach that evaluates cortical activation through hemodynamic response estimation. This study compared the FC of bilateral motor and prefrontal cortices between responders and nonresponders to the conditioned pain modulation test (CPM-test) induced by hand immersion in cold water (0–1°C). We included 37 women with fibromyalgia according to the American College of Rheumatology diagnoses criteria (n = 23 responders to CPM-test and n = 14 nonresponders). After the adjustment for multiple comparisons, we found that nonresponders relative to responders showed higher FC between the left motor cortex (LMC) with the left prefrontal cortex (LPFC) and right prefrontal cortex (RPFC). The psychiatric diagnoses were also positively associated with a higher FC in LMC–LPFC and LMC–RPFC. These results indicate that the increased connectivity between the left motor and bilateral prefrontal cortex might be a neural marker of DPMS dysfunction and an intermediate in the interplay between fibromyalgia and psychiatric disorders.

## Introduction

Previous studies have presented evidence of prefrontal cortex (PFC) inhibitory dysfunction in chronic pain syndromes, including fibromyalgia [10]. Furthermore, a growing pool of evidence supports that motor function plays a pivotal role in pain syndrome outcomes [13]. Grey matter loss indicates cortical anatomical abnormalities in fibromyalgia and other pain conditions [35, 37, 38]. Reversal of gray matter loss in the PFC and motor cortex (MC) occurred following therapeutic interventions in chronic pain patients [50, 51, 58, 59]. These findings suggest that chronic pain’s underlying pathology is not neurodegenerative and depends structurally and functionally on these cortical areas, which are pivotal for integrating the descending pain modulatory system (DPMS) [22, 45, 66].

DPMS is a set of neural networks that encompasses subcortical structures-e.g., the periaqueductal substance (PAG), rostral medial medulla, nucleus accumbens, mesolimbic reward circuit, and amygdala-, as well as cortical areas-e.g., the MC, the PFC, the primary (S1) and secondary somatosensory cortex (S2), and the cingulate [20, 46]. Among these structures, PFC and the limbic system play a significant role in encoding pain’s emotional and contextual components [8, 36, 67]. In contrast, the MC is a significant top-down inhibitory effector, representing a promising area for neuromodulatory intervention [21, 69]. The function of DPMS in fibromyalgia may be assessed by the conditioned pain modulation test (CPM-test) [10], in which higher dysfunction levels correlate with greater fibromyalgia symptom severity and functional disability [60].

In addition to anatomical changes, interest in understanding the functional connectivity (FC) among these cortical areas is rapidly growing. Functional near-infrared spectroscopy (fNIRS) is an imaging method with an excellent temporal resolution that detects changes in oxygenated (HbO) and deoxygenated hemoglobin levels due to the transient dynamics of neurovascular coupling elicited by neuronal activation, a phenomenon that constitutes the hemodynamic response [3, 52, 56, 69]. Regarding fibromyalgia, studies showed intrinsic connectivity abnormalities—when assessed by resting-state functional connectivity—in the DPMS [9, 68], default mode network [43], executive attention network [42], cognitive control network [34], the insular and cingulate cortices [29], and sensorimotor network [17]. FC uncovered different system-perturbation patterns between groups when analyzed during tasks, after the application of stimuli or when deployed for long-term comparisons between pre-and post-therapeutic interventions, also having a predictive value of patient’s treatment response [55, 18, 33].

We conducted this study to test whether FC by fNIRS in fibromyalgia patients might constitute a biomarker of the DPMS dysfunction assessed by the CPM-test, evaluated according to changes in the Numerical Pain Scale (NPS) (0–10 points) during the immersion of the dominant hand in cold water [60]. Therefore, we have compared the FC of bilateral MC and PFC between groups of CPM-test responders and nonresponders in female fibromyalgia patients.

## 2. Materials and Methods

### 2.1. Design overview, settings, and participants

This study’s protocol was approved by the Institutional Review Board (IRB) of the Hospital de Clinicas de Porto Alegre (HCPA), Brazil, and registered in the Certificate of Presentation of Ethical Appreciation (CAAE registry n° 20170330). We obtained written informed consent from all participants before their inclusion. The study enrollment period ranged from January 2018 to December 2019.

### 2.2. Recruitment and inclusion and exclusion criteria

We included 44 right-handed literate adult females aged 30 to 65 years with fibromyalgia seen in the outpatients’ chronic pain wards of the HCPA and digital media publicity. The diagnosis of fibromyalgia was made according to the American College of Rheumatology (2010–2016) guidelines and confirmed by physicians with more than ten years of experience in chronic pain care. This study included only women due to the higher prevalence of fibromyalgia in women than men [54]. We included only patients with NPS scores greater than or equal to six points. Patients were excluded if they had the following medical conditions: rheumatoid arthritis; systemic lupus erythematosus; or another autoimmune, neurologic, or oncologic disease. Those with current uncompensated clinical disease (e.g., ischemic heart disease, chronic kidney disease, or hepatic disease) or using cannabis or recreational psychotropic drugs in the last six months were similarly excluded.

### 2.3. Sample size estimation

The post-hoc sample-size calculation revealed that a sample of 37 patients would ensure detection of effect size (f2) of 0.25 during multiple regression analysis, allowing three predictors based on the consideration of type I and II errors 0.05 and 0.20, respectively [61].

### 2.4. Dependent and independent variables of primary interest

Ipsi- and contralateral FC between the MC and PFC were the dependent variables. The main factor of interest was the function of DPMS evaluated by the change in the NPS score during the CPM-test.

### 2.5. Instruments and assessments

#### 2.5.1. Functional near-infrared spectroscopy assessment

Cortical activation was evaluated by fNIRS using a NIRx® continuous waveform NirScout® near-infrared spectroscopy device (NIRx Medical Technologies, Glen Head, NY, USA) with a scan rate of 15 Hz and dual-wavelength light-emitting diode sources (760 and 850 nm). We used four sources and 16 detectors spaced about 3 cm apart and placed over the scalp using caps provided by EASYCAP®, with 16 channels in total. Probe localization was established using the international 10/10 electroencephalography system. In our montage, we placed sources in the F3, F4, C3, and C4 locations and detectors in the AF3, F5, FC3, F1, C5, CP3, C1, AF4, F6, FC4, F2, C6, CP4, and C2 locations, respectively. Recordings were made using the NirsStar® version 14.2 software program (NIRx Medical Technologies, Glen Head, NY, USA) and were compiled during a single session for each patient. After adjusting the NIRS cap in the scalp, a black cover was placed over it to reduce the degree of environmental light disturbance. Source-detection calibration and recording-checked signal quality were only confirmed if the calibration returned an assessment of “excellent” quality in at least 14 out of 16 channels and if the other two channels were at least “acceptable” on a qualitative scale based on gain, amplitude, coefficient of variation of noise, and dark noise. The resting-state activity was recorded for seven minutes with patients in a sitting position, during which time they were asked to think about nothing while fixating their gaze on the front wall.

##### 2.4.1.1. Functional connectivity analysis

Imaging data were preprocessed using the Brain AnalyzIR® [52] software on the MATLAB (MathWorks, Natick, MA, USA) platform. For the assessment of functional connectivity values, raw data were downsampled to 1 Hz to adequately address the high level of temporal autocorrelation in fNIRS signals. They were subsequently converted to optical density and then to oxyhemoglobin concentration variation (HbO) using the Beer-Lambert law’s modified version [30].

Data were treated through an autoregressive prewhitening model to correct structured noisy and serially correlated error effects (e.g., physiological noise) together with iterative reweighting through a robust regression method addressing outliers (e.g., motion artifacts) with no use of any band/high/low filtering [4, 28]. This method is appropriate for the necessary treatment of serially correlated errors, colored noise, and motion artifacts, dispensing other forms of data processing [4, 5]. In this study, the HbO signal was included in the analysis based on previous reports, which have indicated that this is the most sensitive variable by which to estimate cortical activity via making inferences concerning neurovascular coupling based on hemodynamic response [20, 27, 63, 72,]. Finally, Pearson correlation values were computed for all possible pairs of channels across the time series, and the resulting sample correlation coefficients underwent a Fisher Z-transformation. The Z-values were then averaged for each region of interest (ROI) in each subject (four ROIs × 37 subjects) [53].

#### 2.4.2. Conditioned pain modulation test

A thermode (30 × 30 mm) was attached to the skin on the mid-forearm ventral aspect, and the temperature was set at 32ºC. The thermode was heated at a rate of 1°C per second up to 52°C, where the temperature was set to drop again thereafter. The CPM-test assessment encompassed three stages: (i) we performed quantitative sensory testing (QST), and subjects reported when the temperature reached a level of pain consistent with an NPS score of six points (i.e., “6/10 score”) (T0). We replicated this assessment three times and averaged the temperatures to determine the necessary value to achieve an NPS 6/10 score. All three replicates were performed with an interstimulus interval of 40 seconds, and the position of the thermode was slightly altered between trials. (ii) Five minutes after assessing stage I, patients immersed their right hands up to their wrists into the water at a temperature of between 0°C and 1°C for one minute. After 30 seconds of hand immersion, the QST was reapplied on the area of the thermode used in stage I. (iii) The CPM-test score was calculated according to the change in the NPS score retrieved during the cold-water immersion (QST + CPM-test) at the temperature set as 6/10 on the NPS during the initial trial (T0). Patients were considered responders to the CPM-test if they showed a difference of less than zero and nonresponders if the difference was greater than or equal to zero [7]. It is essential to realize that the CPM-test’s negative values suggest a higher effect of heterotopic stimulus inhibiting the test stimulus (i.e., “pain inhibits pain”)—in other words, a better function of the DPMS.

#### 2.4.3. Pain measures, psychological assessments, sleep quality, and sociodemographic characteristics

We used the Fibromyalgia Impact Questionnaire (FIQ) [41] to evaluate the impact of life quality symptoms. VAS Analogue Scale assessed the severity of scores in most days of the last three months ranged (VAS, zero to 100mm). The Pain Catastrophizing Scale (PCS) [57] was used to assess the identification of an individual’s pain catastrophizing into three dimensions: rumination, magnification, and helplessness. Beck Depression Inventory-II [23] was used to evaluate depressive symptoms, and Central Sensitization Inventory [12] was used to measure symptoms related to central sensitization syndrome. Over the last month, the sleep quality and sleep patterns were evaluated by the Pittsburgh Sleep Quality Index (PSIQI) [6].

We used a standardized query to assess demographic data and medical comorbidities. We requested to provide information about their age, sex, years of education, and lifestyle habits. Patients also provided information about their health status, including clinical and psychiatric diagnoses, the latter assessed by the Mini-international Neuropsychiatric Interview (M.I.N.I.) [1]. A specific questionnaire evaluates all medications used and their daily doses (e.g., antidepressants, anticonvulsant, hypnotic, analgesics non-opioid and opioid, etc.). We calculated the opioid use by mean morphine-equivalent dose (MED) per day [70]. They were classified with minimal opioid use or no use if the mean daily MED was less than 5 mg or if self-reported opioid use was less than twice a week in the previous 28 days. If they self-reported a mean daily MED equal to or greater than 5 mg for most days for the last three months, they were classified with regular opioid user. Non-opioid analgesic use was defined by analgesic use per week on most days of the previous month. For data analysis, the analgesic use was dichotomized in those used analgesics at least four or fewer days per week or the use for more than four days per week. We adopted this approach because chronic pain patients can change their rescue analgesic use each week depending on pain levels.

Among the efforts to reduce potential sources of bias, all assessments were conducted by researchers with the vast clinical expertise to treat outpatients in a pain clinic, and the diagnostics were done according to the pre-specified criteria described above. Two evaluators with specific training were responsible for all assessments and to apply the standardized protocol to assess the CPM-test and the evaluation using the fNIRS.

### 2.5. Statistical Analysis

Chi-squared, Fisher’s exact tests, and t-tests for independent samples were used to compare categorical and continuous variables between groups. We used the Shapiro-Wilk test to test for normality. The FC was compared by t-test for independent samples in univariate analysis. A multivariate covariance analysis (MANCOVA) model was used to compare FC Z-values among four ROIs taken in pairs (ROIs: left MC, right MC; left PFC, right PFC) between responders and nonresponders adjusted by the following factors: opioid analgesic use (Minimal/no use, or regular opioid use), non-opioid analgesic use, and psychiatric disorder diagnosis (according to the MINI Yes/No). To identify the source of significant differences was used the Bonferroni’s Multiple Comparison Test. Statistical significance was set to a p-value of 0.05, 2-tailed. Statistical analysis was conducted in SPSS software version 22.0 (SPSS, Chicago, IL).

## 3. Results

### 3.1. Patient characteristics

We enrolled 43 subjects, from which six were excluded, three for missing data, and three due to problems in the fNIRS measures. The final sample was composed of 37 participants (23 responders and 14 nonresponders to CPM-test). The characteristics of the sample and comparative analyses between responders and nonresponders are presented in Table 1. We did not find statistical differences between responders and nonresponders in sociodemographic characteristics, psychological measures, sleep quality, psychotropic medications, and analgesic use. However, nonresponders, compared to responders, presented higher scores in the Visual Analogue Scale (p=0.02) and higher psychiatric disorder prevalence (p=0.03).

**Table 1.**
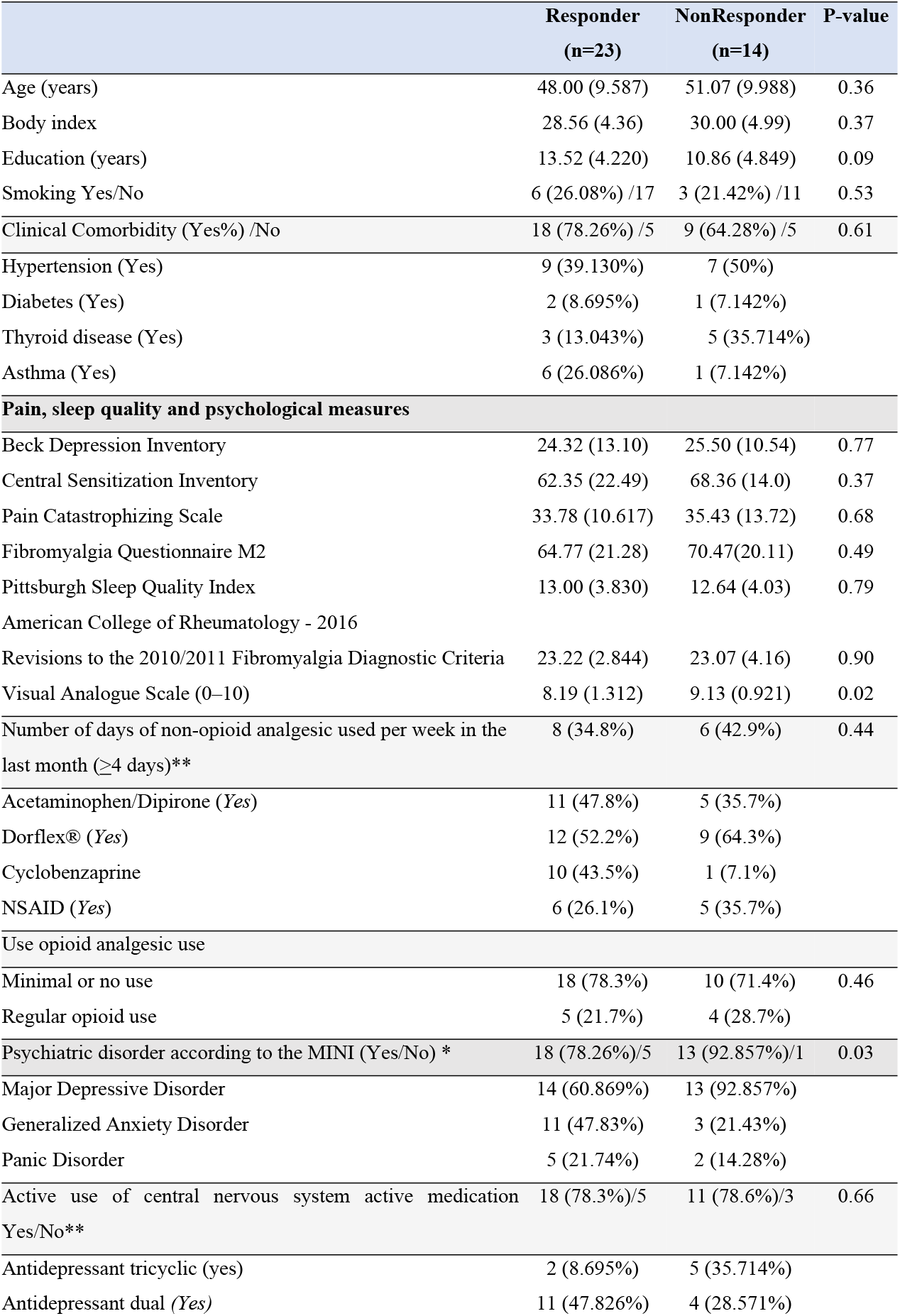
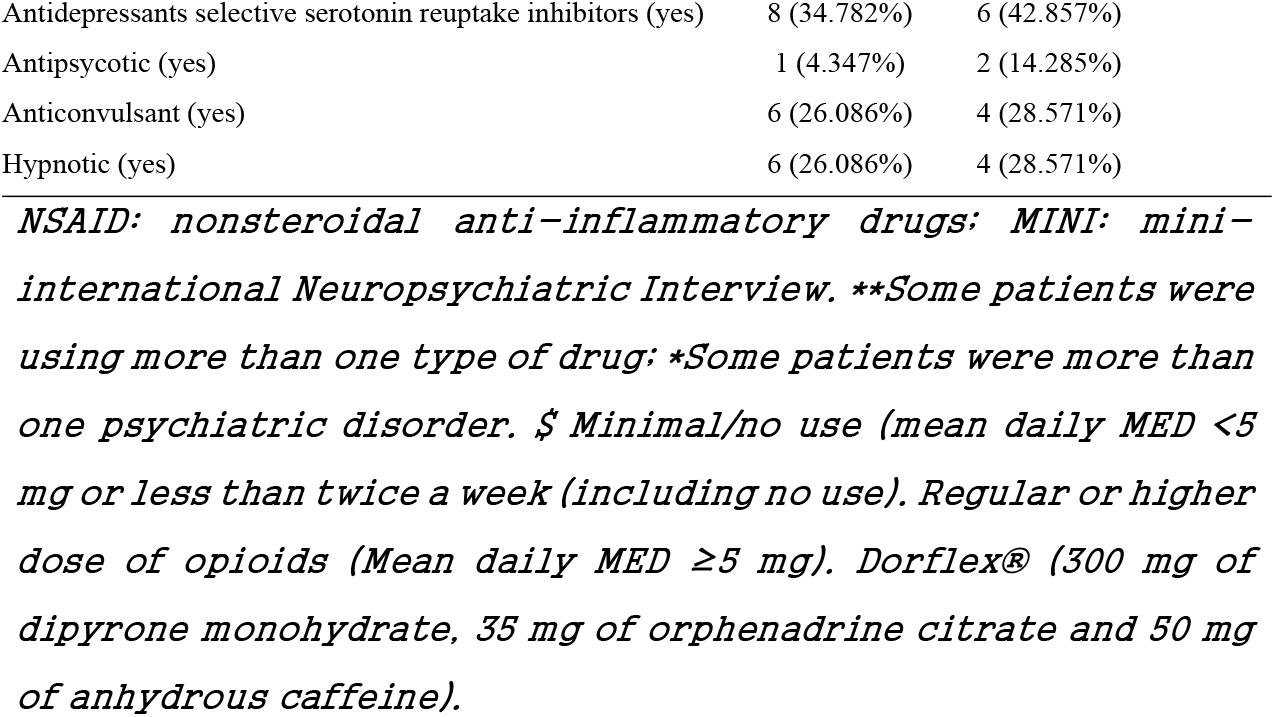
Demographic and clinical characteristics of the study sample. Values are given as the mean (SD) or frequency (n=37).

### 3.2. Univariate analysis according to the spectrum of responders and nonresponders to CPM-test in the FC

According to the spectrum to responders and nonresponders to CPM-test, the mean (Standard Deviation) of cortical FC is presented in Table 2. Nonresponders to CPM-test showed higher FC between the left MC and the left PFC (t=-2.476, p=0.01) compared to responders. Also, nonresponders presented higher FC levels between the left MC and the right PFC (t=-2.363, p=0.02).

**Table 2.** Relationship of the outcomes and the main interest factor according to the spectrum of responders and non-responders to CPM-test. Data are presented as the mean (standard deviation) (n=37).

| Region of interest connectivity | Responders | Nonresponders | t | p-value |
| --- | --- | --- | --- | --- |
| Left prefrontal cortex – Left motor cortex | 0.607 (0.165) | 0.738 (0.138) | -2.476 | 0.01 |
| Left prefrontal cortex – Right prefrontal cortex | 0.728 (0.189) | 0.778 (0.133) | -0.861 | 0.39 |
| Left prefrontal cortex – Right motor cortex | 0.521 (0.163) | 0.608 (0.085) | -1.831 | 0.07 |
| Left motor cortex – Right prefrontal cortex | 0.524 (0.165) | 0.648 (0.131) | -2.363 | 0.02 |
| Left motor cortex – Right motor cortex | 0.518 (0.224) | 0.610 (0.149) | -1.370 | 0.17 |
| Right prefrontal cortex – Right motor cortex | 0.544 (0.199) | 0.617 (0.133) | -1.214 | 0.23 |

**Table 3.**
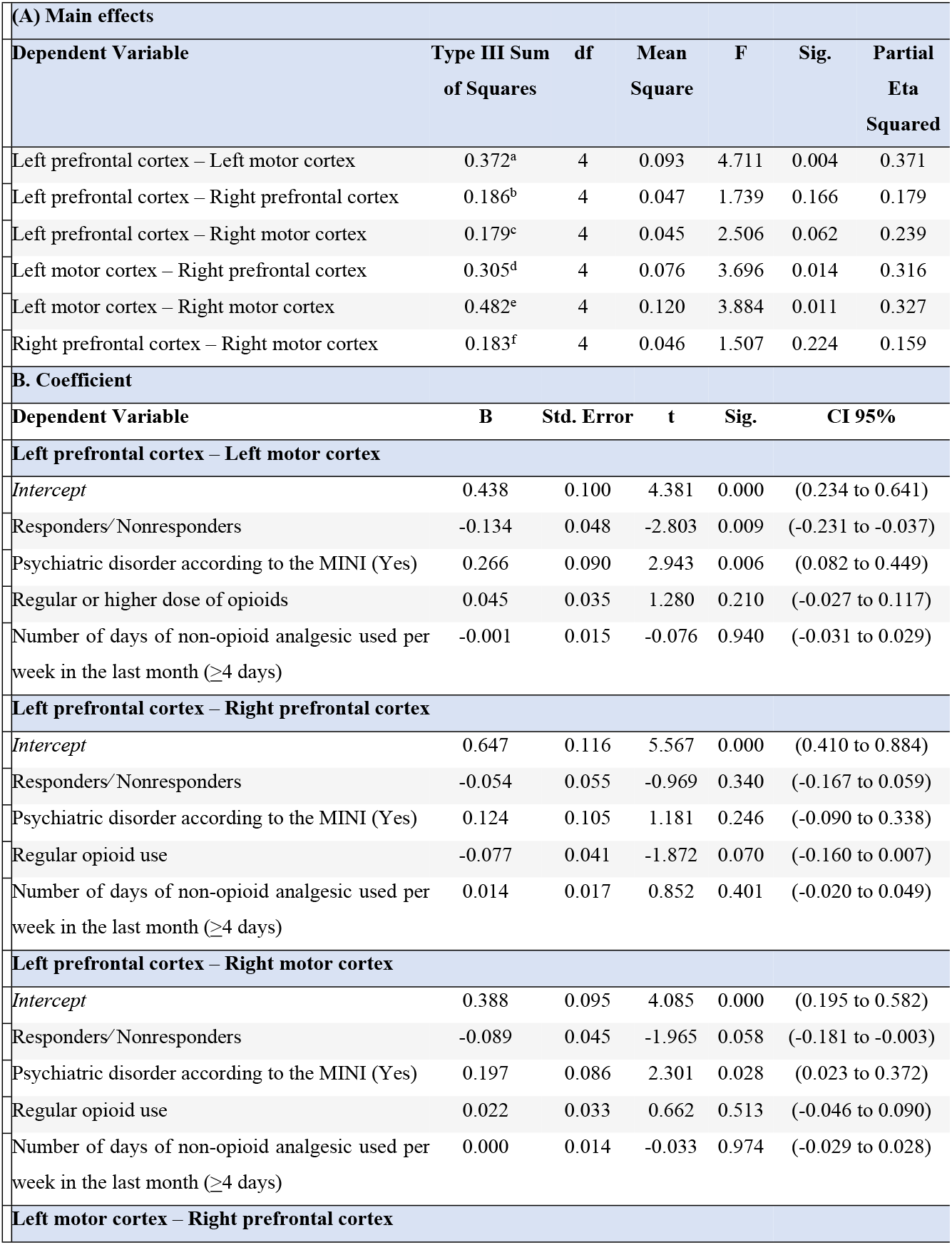
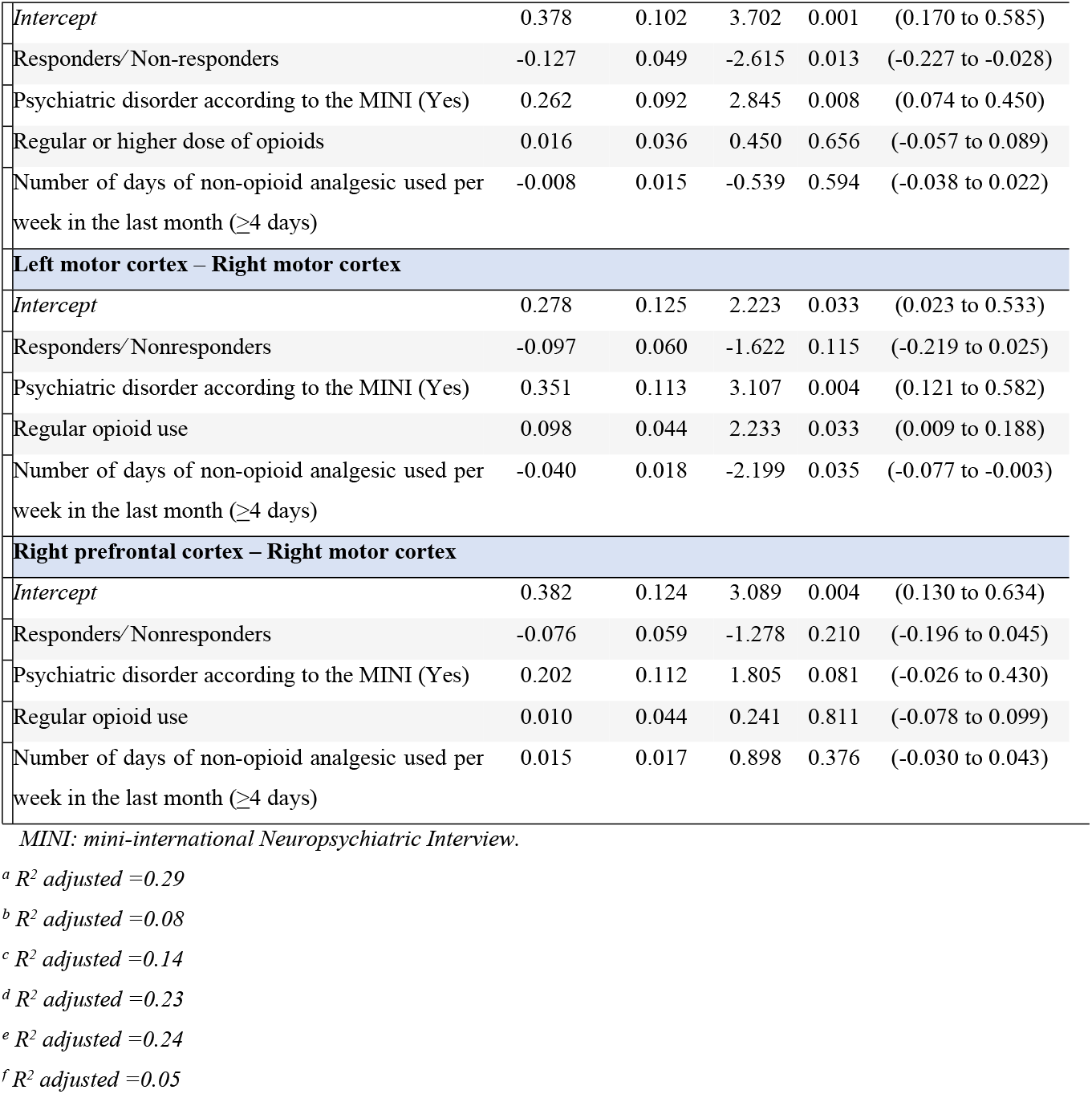
MANCOVA analysis of the relationship between the FC according to the spectrum of responders or non-responders according to change in NPS (0-10) during the CPM-test (n=37).

| (A) Main effects |  |  |  |  |  |  |
| --- | --- | --- | --- | --- | --- | --- |
| Dependent Variable | Type III Sum of Squares | df | Mean Square | F | Sig. | Partial Eta Squared |
| Left prefrontal cortex – Left motor cortex | 0.372<sup>a</sup> | 4 | 0.093 | 4.711 | 0.004 | 0.371 |
| Left prefrontal cortex – Right prefrontal cortex | 0.186<sup>b</sup> | 4 | 0.047 | 1.739 | 0.166 | 0.179 |
| Left prefrontal cortex – Right motor cortex | 0.179<sup>c</sup> | 4 | 0.045 | 2.506 | 0.062 | 0.239 |
| Left motor cortex – Right prefrontal cortex | 0.305<sup>d</sup> | 4 | 0.076 | 3.696 | 0.014 | 0.316 |
| Left motor cortex – Right motor cortex | 0.482<sup>e</sup> | 4 | 0.120 | 3.884 | 0.011 | 0.327 |
| Right prefrontal cortex – Right motor cortex | 0.183<sup>f</sup> | 4 | 0.046 | 1.507 | 0.224 | 0.159 |
| B. Coefficient |  |  |  |  |  |  |
| Dependent Variable | B | Std. Error | t | Sig. | CI 95% |  |
| Left prefrontal cortex – Left motor cortex |  |  |  |  |  |  |
| Intercept | 0.438 | 0.100 | 4.381 | 0.000 | (0.234 to 0.641) |  |
| Responders/Nonresponders | -0.134 | 0.048 | -2.803 | 0.009 | (-0.231 to -0.037) |  |
| Psychiatric disorder according to the MINI (Yes) | 0.266 | 0.090 | 2.943 | 0.006 | (0.082 to 0.449) |  |
| Regular or higher dose of opioids | 0.045 | 0.035 | 1.280 | 0.210 | (-0.027 to 0.117) |  |
| Number of days of non-opioid analgesic used per week in the last month (≥4 days) | -0.001 | 0.015 | -0.076 | 0.940 | (-0.031 to 0.029) |  |
| Left prefrontal cortex – Right prefrontal cortex |  |  |  |  |  |  |
| Intercept | 0.647 | 0.116 | 5.567 | 0.000 | (0.410 to 0.884) |  |
| Responders/Nonresponders | -0.054 | 0.055 | -0.969 | 0.340 | (-0.167 to 0.059) |  |
| Psychiatric disorder according to the MINI (Yes) | 0.124 | 0.105 | 1.181 | 0.246 | (-0.090 to 0.338) |  |
| Regular opioid use | -0.077 | 0.041 | -1.872 | 0.070 | (-0.160 to 0.007) |  |
| Number of days of non-opioid analgesic used per week in the last month (≥4 days) | 0.014 | 0.017 | 0.852 | 0.401 | (-0.020 to 0.049) |  |
| Left prefrontal cortex – Right motor cortex |  |  |  |  |  |  |
| Intercept | 0.388 | 0.095 | 4.085 | 0.000 | (0.195 to 0.582) |  |
| Responders/Nonresponders | -0.089 | 0.045 | -1.965 | 0.058 | (-0.181 to -0.003) |  |
| Psychiatric disorder according to the MINI (Yes) | 0.197 | 0.086 | 2.301 | 0.028 | (0.023 to 0.372) |  |
| Regular opioid use | 0.022 | 0.033 | 0.662 | 0.513 | (-0.046 to 0.090) |  |
| Number of days of non-opioid analgesic used per week in the last month (≥4 days) | 0.000 | 0.014 | -0.033 | 0.974 | (-0.029 to 0.028) |  |
| Left motor cortex – Right prefrontal cortex |  |  |  |  |  |  |

|  |  |  |  |  |  |
| --- | --- | --- | --- | --- | --- |
| *Intercept* | 0.378 | 0.102 | 3.702 | 0.001 | (0.170 to 0.585) |
| Responders/Non-responders | -0.127 | 0.049 | -2.615 | 0.013 | (-0.227 to -0.028) |
| Psychiatric disorder according to the MINI (Yes) | 0.262 | 0.092 | 2.845 | 0.008 | (0.074 to 0.450) |
| Regular or higher dose of opioids | 0.016 | 0.036 | 0.450 | 0.656 | (-0.057 to 0.089) |
| Number of days of non-opioid analgesic used per week in the last month ($\geq 4$ days) | -0.008 | 0.015 | -0.539 | 0.594 | (-0.038 to 0.022) |
| **Left motor cortex – Right motor cortex** |  |  |  |  |  |
| *Intercept* | 0.278 | 0.125 | 2.223 | 0.033 | (0.023 to 0.533) |
| Responders/Nonresponders | -0.097 | 0.060 | -1.622 | 0.115 | (-0.219 to 0.025) |
| Psychiatric disorder according to the MINI (Yes) | 0.351 | 0.113 | 3.107 | 0.004 | (0.121 to 0.582) |
| Regular opioid use | 0.098 | 0.044 | 2.233 | 0.033 | (0.009 to 0.188) |
| Number of days of non-opioid analgesic used per week in the last month ($\geq 4$ days) | -0.040 | 0.018 | -2.199 | 0.035 | (-0.077 to -0.003) |
| **Right prefrontal cortex – Right motor cortex** |  |  |  |  |  |
| *Intercept* | 0.382 | 0.124 | 3.089 | 0.004 | (0.130 to 0.634) |
| Responders/Nonresponders | -0.076 | 0.059 | -1.278 | 0.210 | (-0.196 to 0.045) |
| Psychiatric disorder according to the MINI (Yes) | 0.202 | 0.112 | 1.805 | 0.081 | (-0.026 to 0.430) |
| Regular opioid use | 0.010 | 0.044 | 0.241 | 0.811 | (-0.078 to 0.099) |
| Number of days of non-opioid analgesic used per week in the last month ($\geq 4$ days) | 0.015 | 0.017 | 0.898 | 0.376 | (-0.030 to 0.043) |

*MINI: mini-international Neuropsychiatric Interview.*
<sup>a</sup> $R^2$ adjusted =0.29
<sup>b</sup> $R^2$ adjusted =0.08
<sup>c</sup> $R^2$ adjusted =0.14
<sup>d</sup> $R^2$ adjusted =0.23
<sup>e</sup> $R^2$ adjusted =0.24
<sup>f</sup> $R^2$ adjusted =0.05

### 3.3. Multivariate analysis of the relationship between the FC according to the spectrum of responders and nonresponders to CPM test

The results of the MANCOVA model analysis with dependent variables, the FC between MC and PFC according to the spectrum of responders and nonresponders to the CPM-test, and the number of psychiatric disorders, number of non-opioid analgesics used daily, and number of opioid medications as independent variables, are presented in Table The MANCOVA model using Bonferroni’s Multiple Comparison Test revealed a significant relationship between the responders and nonresponders groups and the outcomes related to FC (Hotelling’s Trace =1.21, F(6)=5.47, P=0.001). This analysis presented a power of 0.98.

In the regression analysis, nonresponders showed higher FC between the left MC and the left and right PFC. The left PFC and the right MC displayed increased FC in nonresponders with a borderline statistical significance (p=0.058). Cortical FC between the left MC and the right MC was significant on the main effect, with no difference between groups when considered covariates in the regression analysis. No group effect was observed between the right PFC and right MC.

Psychiatric disorders were associated with FC between left MC-left PFC, left MC-right PFC, left PFC-right MC, and left MC-right MC. In contrast, the regular opioid and non-opioid analgesic use were associated with increased interhemispheric FC in MC. The map of FC across regions of interest in Z-values upon thermal stimulus is shown in Figure 3.

**Figure 1.**
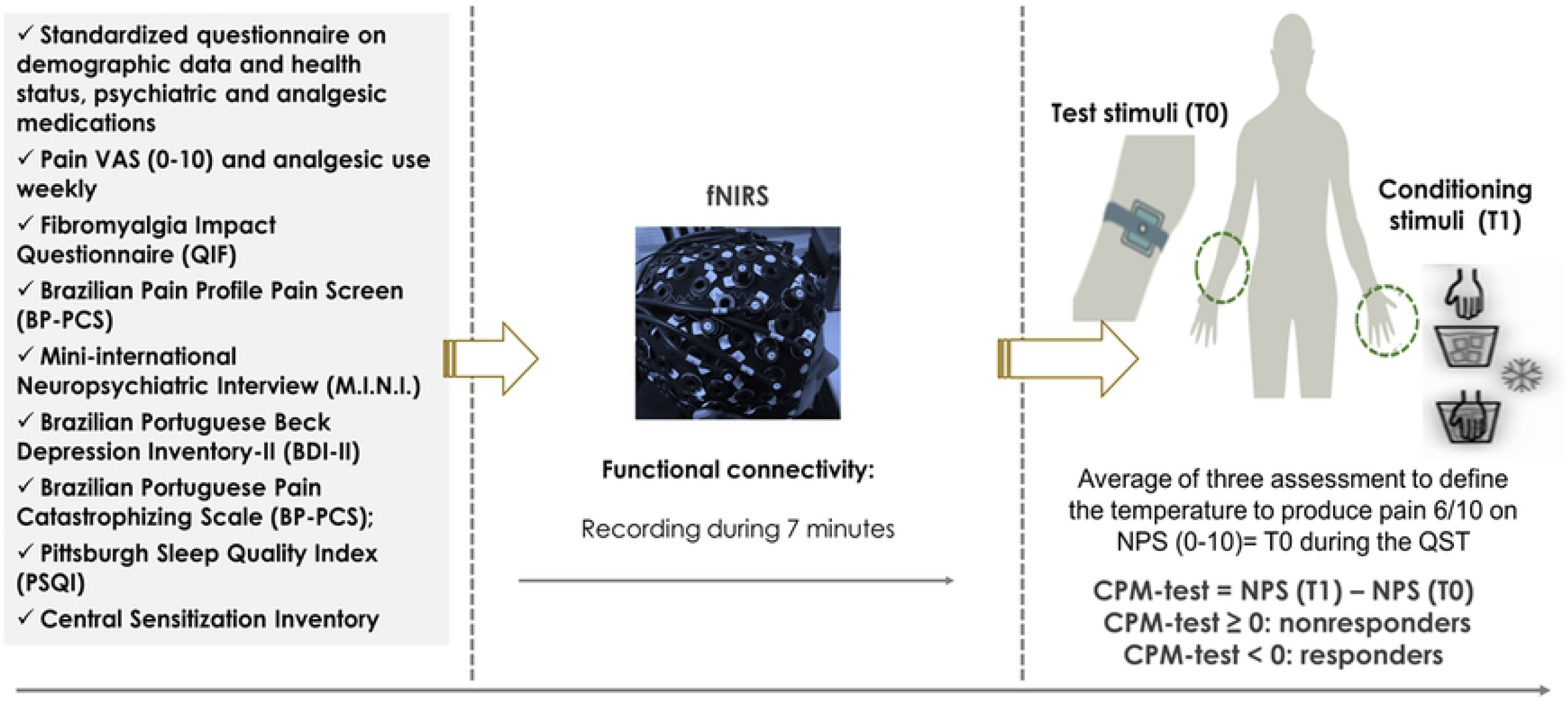
Timeline assessments. CPM-test: condition pain modulation test; fNIRS: functional near-infrared spectroscopy.

**Figure 2.**
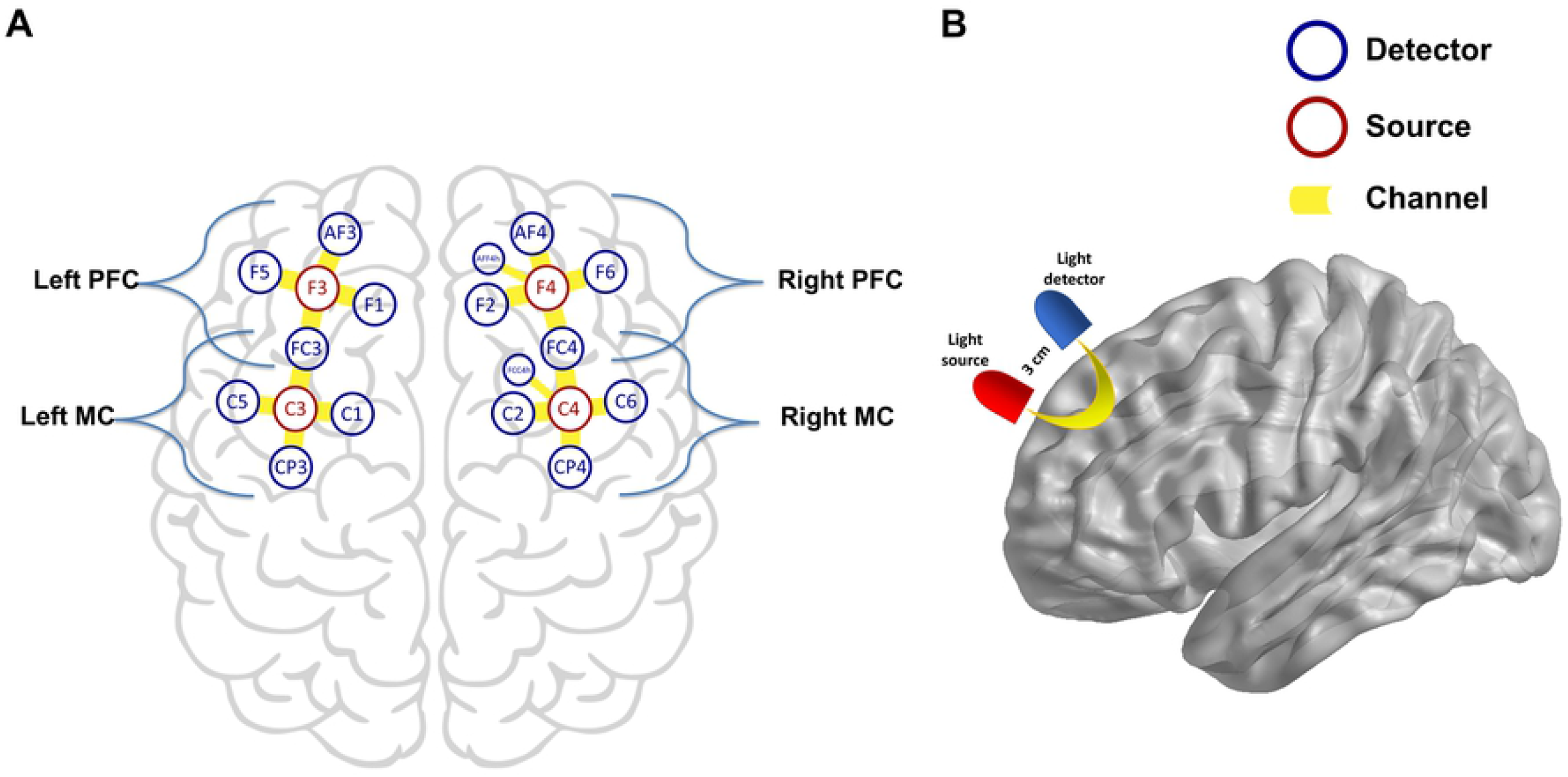
fNIRS montage. **A**. Sources and detectors were arrayed in 10-10 system. AFF3h, FCC3h, AFF4h, FCC4h correspond to short-distance channels. **B**. Illustrative demonstration of near-infrared diffuse reflection path between source and receptor.

**Figure 3.**
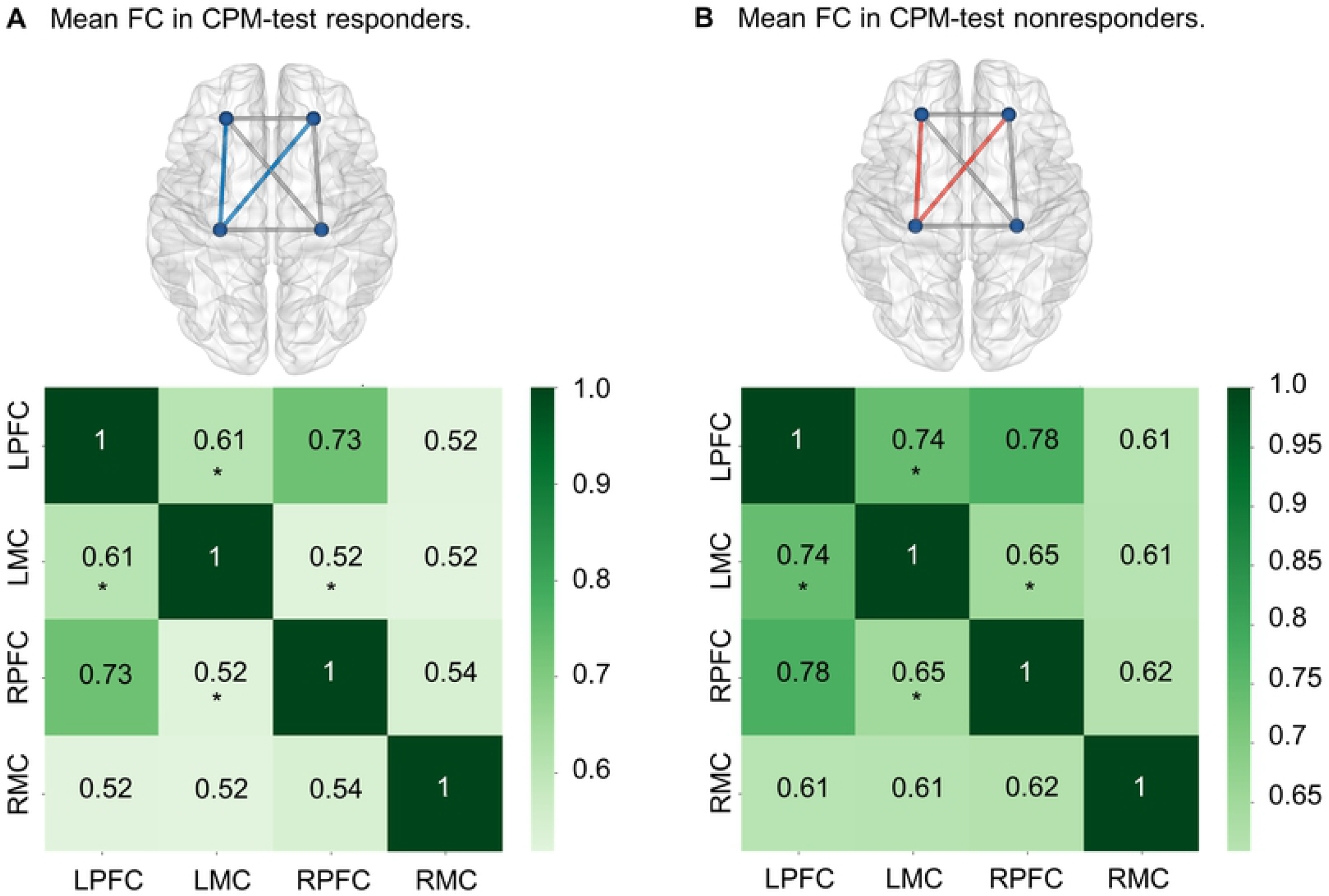
Heatmap visualization of FC across regions of interest in Z-values. Nonresponders represented higher connectivity than responders between LMC and bilateral PFC. **A**. Mean FC by region of interest in CPM-test responders. **B**. Mean FC by region of interest in CPM-test nonresponders. FC: functional connectivity; CPM-test: conditioned pain modulation test. LPFC: Left prefrontal cortex; LMC: Left motor cortex; RPFC: Right prefrontal cortex; RMC: Right motor cortex. *p<0.05.

## 4. Discussion

Our findings indicate that nonresponders have an increased FC of left MC with left and right PFC, with a higher FC between left MC and right MC not attributed to the CPM-test but other covariates (i.e., regular opioid use and psychiatric disorders’ coexistence). Conversely, non-opioid analgesics are negatively related to inter-hemispheric FC between right MC and left MC. Together, these results suggest that the increased FC between left MC with bilateral PFC is a neural correlate linked to the dysfunction of the DPMS. Thus, our results concatenate the activity imbalance between brain regions with the inhibitory deficiency of the DPMS.

Considering that the CPM-test assesses the potency of the DPMS, the increased FC between left MC with right and left PFC is aligned with the perspective that these findings seem to represent a marker of dysfunction in the inhibitory potency of DPMS. According to an earlier study, the hyper inhibition at the cortical level was associated with higher dysfunction of DPMS in fibromyalgia [10]. Thus, these areas’ hyper-connectivity suggests that the FC may be a marker of cortical network reorganization, and it highlights that this cortical restructuring is associated with DPMS dysfunction. These functional changes may also be processes underpinning central sensitization’s pathophysiology, integrating cortical brain networks’ deterioration with the corticospinal pain pathways [60]. Consistent with our findings of increased FC in DPMS dysfunction, earlier fMRI studies found that in patients with chronic pain, including chronic low back pain and fibromyalgia, the increased FC between the insula cortex and default mode network, including the medial PFC, is correlated with pain perception intensity [11, 39, 42]. Furthermore, in previous studies, pharmacological and physical interventions to reduce pain also decreased such connectivity, while maneuvers to increase pain increased the FC [25, 39, 43]. The current results and the previous fMRI studies suggest that enhanced FC between the MC and PFC is critical to sustaining chronic pain and fibromyalgia symptoms.

Interestingly, higher connectivity was found in the primary cortical areas involved in pain processing. The increased FC between left MC with PFC areas may indicate a disruption on the mechanism mediated by inhibitory gamma-aminobutyric acid (GABAergic) on the primary motor cortex interneurons, as indexed by short intracortical inhibition (SICI) [15]. Simultaneously, it suggests an up regulatory phenomenon of intracortical inhibitory networks mediated by GABA receptors, a plausible hypothesis according to an earlier study that found an association of increased transient SICI with the deficiency of the DPMS [10]. Hence, these results build in knowledge on the increased connection between MC and PFC’s neural network associated with DPMS, which may be a compensatory mechanism to counter-regulate the sustained cortical hyperexcitability in chronic pain. The concept of a physiological adaptation response following a prolonged system demand supports this hypothesis and may explain the dysfunctional remapping of related system components, which has been observed in other contexts with autonomic, metabolic, and inflammatory systems [24].

The MC inhibits pain by top-down efferences on the DPMS. Thus, these connections may subsidize the MC as a therapeutic target for the transcranial neuromodulation used to treat pain. In the same way, the asymmetric functions of the PFC brain hemispheres can explain the benefits of using excitatory stimulation on the PFC to treat chronic pain or depression, such as the high frequency with repetitive transcranial magnetic stimulation (rTMS) or anodal transcranial direct current stimulation [49, 26]. In contrast, rTMS with low frequency has been used to treat depression when the rTMS is applied to the right PFC. Furthermore, dorsolateral PFC (DLPFC) inhibitory effects on the ipsilateral primary motor cortex (M1) is higher in the left than in the right hemisphere [71], even though right DLPFC has inhibitory effects on the left M1 [44]. Right-handers present stronger effects from the left-to-right MC, which seems to involve excitatory and inhibitory inter-hemispheric activity [73], supporting the idea of an increased activity tone in right-handers’ left M1. Our results regarding connectivity between PFC and the left MC may be influenced by the hemispheric dominance when considering that our sample was composed solely of right-hander patients.

The analgesic effect of MC is correlated with the release of endogenous opioids in the anterior cingulate cortex (ACC), insula, and periaqueductal gray matter (PAG) [74]. The anterior MC receives projections from the spinothalamic system transmitting nociceptive information to the cortex, and its activation can be achieved by global pain network modulation. Studies of functional neuroimaging have shown that MC activates supraspinal areas involved in pain perception and emotional appraisal. In a preclinical experiment of neuropathic pain, the MC stimulation reversed the central and peripheral pain [40], activating the limbic system and PAG and inhibiting the thalamic nuclei and spinal nociceptive neurons [47].

The PFC role in pain processing is integrated with other brain areas, including primary and secondary somatosensory cortices, insular cortex (IC), ACC, and thalamus [2]. The discriminative pain intensity involves connections of the ventrolateral pathways, from the insular cortex to the PFC. Activation of the ACC and medial-PFC (m-PFC) is associated with increased activity on the PAG [48], which is the main control center for descending pain modulation. The PAG originates principally from the PFC cortical projections to the PAG. In contrast, the ventrolateral-PAG is linked to brain regions associated with descending pain modulation, including the ACC, upper pons/medulla [14]. There is a decrease in the glutamate and reductions in the gray matter density in chronic pain in the PFC. Chronic pain may result from disruption of this descending analgesic circuitry that stems from PFC deactivation. Thus, the FC enhancement between PFC and MC in the ipsi- and contralateral hemispheres points towards critical dysfunction on the nociceptive circuitry in fibromyalgia.

Together, these results give insights into generating testable hypotheses regarding improving and optimizing therapy. These results might be applicable for top-down neuromodulation using the tDCS and TMS. These techniques target theM1 and the DLPFC [74]. Variation in FC between the left MC and the PFC in a DPMS-dependent magnitude is a result that highlights the criticality of those cortical areas in pain neuromodulation. These findings also shed light on the mechanism of action of tDCS as a treatment for chronic pain by hinting at the FC abnormalities as potential direct targets and interest variables during follow-up.

It has long been suggested that optically recorded hemodynamic response is not necessarily equivalent to the pyramidal cell activity. Given that cortical layers are constituted by dense interconnections of pyramidal neurons and inhibitory interneurons, it is impossible to differentiate - by estimation of hemodynamic response alone - which information is being outputted by presynaptic neuronal activity [16, 31, 32, 64]. Hence, the higher intensity in a particular ROI implicates a higher presynaptic neural activity independently of its activity quality (e.g., excitatory or inhibitory synaptic signaling). Nevertheless, previous studies on knee osteoarthritis chronic pain patients have found the left MC to display higher corticospinal excitability (assessed by motor-evoked potential) and disinhibited state (attested by shortened cortical silent period), which supports the findings of higher intracortical disinhibition in dysfunctional DPMS attested by non-responsiveness to the CPM test in other studies [65]. Central sensitization in chronic pain is usually linked with insufficient descending inhibition signaling and amplification of afferent signals. These neuroplastic changes are involved in allodynia, hyperalgesia, expanded receptive field, prolonged electrophysiological discharge, and neuronal hyperexcitability [19, 62]. Remarkably, this study’s increase in cortical connectivity was associated with a lack of DPMS adaptability, which may suggest an increase in excitatory synaptic signaling and a disinhibited state when taking the above mentioned into account.

The design of this study prevents establishing a causal nexus. This way is required prospective; notwithstanding this limitation, the present work indicates that the cortical FC of non-responder patients is significantly different from responders. We also observed that FC changes correlated with higher opioids use and psychiatric disorders. Although such correlation does not prove direct causality, it is compatible with a causal model in which higher doses of opioids might cause cumulative anatomical–functional changes in pain pathways. In contrast to opioids, the non-opioid analgesic used was inversely correlated with the connectivity, indicating that they might be more controlled.

In conclusion, these results give insights into understanding the reorganization of neural networks in the main areas involved in pain processing. These results indicate that the hyperconnectivity between the motor and prefrontal cortex might be a neural marker of the DPMS dysfunction and an intermediate in the interplay between fibromyalgia and psychiatric disorders.

## DISCLOSURES

### Research funding for Brazilian agencies

i) Committee for the Development of Higher Education Personnel—CAPES (grant no. 2018 to P. V. and R. L. for doctorate scholarship). (ii) National Council for Scientific and Technological Development - CNPq (grant nos. 302688/2017-0 and 420826/2018-1 to W. C.). (iii) Postgraduate Research Group at the Hospital de Clínicas de Porto Alegre (grant no. 20170330). Brazilian Innovation Agency (FINEP [Financiadora de Estudos e Projetos]) (process number 1245/13; grant nos. Fundação de Amparo à Pesquisa do Estado do Rio Grande do Sul (FAPERGS)-17/2551-0001 087-6 and 17/2551-0001 476-6).

W. C. agrees to be accountable for all aspects of the work, ensuring that questions related to the accuracy or integrity of any part of the work are appropriately investigated and resolved.

### Contributorship

*Álvaro de Oliveira Franco, Iraci Lucena da S. Torres, Felipe Fregni, and Wolnei Caumo* were involved in the study concept and design, acquisition of data or analysis, and interpretation of the data. *Álvaro de Oliveira Franco, Camila Fernandes, Paulo Vicunha, and Maria Adelia Aratanha* were involved in data acquiring, analysis or interpretation. *Álvaro de Oliveira Franco, Felipe Fregni, and Wolnei Caumo* contributed to drafting/revising the manuscript for relevant intellectual content. All authors contributed to the approval of the final version to be published.

## Acknowledgments of analysis support

MAA is employed by NIRx Medizintechnik GmbH. All other authors disclose no potential conflicts of interest for the research, authorship, and publication of this article.

